# Comparison of Software for Prediction of Fraction Absorbed and Unbound in Humans

**DOI:** 10.1101/2025.09.07.674727

**Authors:** Urban Fagerholm

## Abstract

The main objective of this study was to evaluate and compare the performance of 5 PK software, ANDROMEDA by Prosilico 2.0, and 4 free, web-based prediction tools, PkCSM, Swiss ADME, ADMETlab 3.0 and DruMAP 2.0, for predictions of fraction absorbed (f_a_) and unbound (f_u_) in humans. Sets with compounds with available and undisclosed estimates were selected (n=140). The risk that test compounds have been used in training sets for model building (and thereby, influenced and exaggerated the predictive performances) was minimized. At least, these were not included in training sets for ANDROMEDA. One set consisted of compounds that have not been marketed and for which pharmacokinetic information has not been publically disclosed. Quantitative and qualitative evaluations and comparisons were done. For both f_a_ and f_u_, ANDROMEDA was clearly more accurate and balanced than the others, with higher Q^2^ (0.69 vs 0.35 for f_a_; 0.94 *vs* 0.62-0.76 for f_u_), lower mean errors (15 % *vs* 28 %; 2.3- *vs* 4.1- to 36-fold), lower maximum errors (54 % *vs* 92 %; 10- *vs* 30- to 524-fold), more correct predicted classes (70-77 % *vs* 13-54 %), no failed, inconclusive or poor predictions (as found for 3 of the other software), wider application range, and minimal skewness at low values. It had intercepts on f_a_- and f_u_-prediction axes that were ca 1/3 and 1/175 to 1/23 compared to those found for the other software, which is of particular importance. Two software, PkCSM and Swiss ADME, were considered inappropriate, whereas ADMETlab took an intermediate performance position. Apparently, DruMAP was second best performing software. Overall, there was poor performance overlap between the software (7-24 %), with many contradictory predictions. Advantages with ANDROMEDA suggest that this is the software of choice for those that desire adequate predictions of f_a_ and f_u_ in humans and estimates of certainty. The findings are of particular interest for the 3R-process.

## Introduction

Several software for prediction of human pharmacokinetics (PK) are available. These include free, web-based products/services. The applicability of such software depends on factors such as coverage and clinical validity of parameters, compound and estimate ranges, prediction accuracy and precision, and security. Software of choice is also based on desired quality standards.

A major obstacle for external validations and performance evaluations of prediction software is lack of transparency and user-knowledge of which compounds that have been used in training sets for model building. If test compounds in evaluations already have been used in training sets, prediction results will be misleading and exaggerated. Other obstacles include few published and peer-reviewed cross-validations and validations using hold-out sets, and limited disclosure of performance statistics. For example, a Q^2^-value could be good/high, and indicate that a model is robust and well-performing, despite skewness and large errors. Major errors could be hidden by the use of RMSE (instead of mean and maximum errors), and the use of % of compounds with <2-fold error could be a way to hide poor performance of a model with estimates ranging from 0 to 100 %. For example, a default value of 40 % would result in a large portion of prediction errors of maximally 2- to 3-fold.

Lack of clinical validity, low accuracy and skewness are highlighted issue with some *in silico* prediction models and software [1]. Since solubility (S) is poorly correlated to *in vivo* dissolution potential (R^2^=0.13) and seems to exaggerate absorption limitations in humans, and intrinsic metabolic clearance (CL_int_) measured with hepatocytes and microsomes correlates poorly with corresponding *in vivo* parameter (R^2^=0.1-0.4) and has relatively narrow range (LOQ of the same magnitude as median CL_int_ for marketed drugs), *in vitro* data based models for these have limited clinical validity and applicability (Q^2^ approximated to ca 0.05-0.15) [1]. They appear more suited for developing drugs and optimizing compounds for an *in vitro* human.

Fraction absorbed (f_a_) and fraction unbound in plasma (f_u_) are essential parameters for prediction of overall PK, exposure profiles and doses. Prediction models for f_a_ and f_u_ often show moderately high Q^2^, but are typically skewed, with extensive overprediction at low values. An example for f_a_ is the *in silico* model by Suenderhauf et al. [2], with a R^2^ of 0.6 and RMSE of 26 %, but an intercept on the predicted f_a_-axis (at 0 % observed f_a_) of ca 50 % f_a_. Many poorly absorbed compounds (f_a_<1-2 %) were predicted to have f_a_ of 80-100 %. Thus, this model (and also others) seems applicable for highly permeable compounds only/mainly (which cannot be evaluated and confirmed by the model). It was approximated that the software ADMET Predictor (by Simulations Plus Inc.) has a Q^2^ of ca 0.25 for f_a_ [1], and with no apparent ability to distinguish between low to moderately high f_a_. On the contrary, Rule of 5 underestimates the absorption potential to a great extent (ca 60 % false negatives and ca 40 % average f_a_ for a selection of rule-violent compounds [3]. Based on these findings and various systematic errors it is difficult to predict f_a_ in humans with high accuracy using *in silico* models, unless a compound is highly permeable and soluble. A large extent (ca 1/3 to 1/2) of modern small compounds has limited permeation and dissolution (incomplete gastrointestinal uptake in humans), and this makes accurate predictions of f_a_ difficult and decision-making uncertain and problematic.

Skewness has also been shown for *in silico*-prediction models and software for f_u_. Yun et al. [4] used 3 different *in silico* f_u_-models/software, ADMET Predictor and models by Watanabe et al. and Ingle et al., and found Q^2^ (or R^2^; unknown how many test compounds that were already used in training sets) of ca 0.4-0.5 and intercepts of ca 4-7 % predicted f_u_ at 1 % observed f_u_ (apparently much greater overpredictions at f_u_<1 %). Many low f_u_ compounds (f_u_<1-3 %) were predicted to have f_u_>50 %. There were also underprediction trends at moderate to high f_u_, with average predicted f_u_ of ca 45-60 % at observed f_u_ of 100 %. Some compounds that do not bind to plasma proteins were predicted to be more than 97 % bound. Thus, the applicability domain of these models are compounds with moderate f_u_. A significant portion (ca 1/3 in 2021) of modern small drugs have f_u_ of maximally 2 % [5]. Skewed and uncertain f_u_-prediction models are anticipated to jeopardize PK-prediction of many drugs under development.

Ultimately, human PK prediction models and software are based on reliable and valid clinical (and/or preclinical) data and produce results with overall high accuracy, wide range and with minimal skewness. We have previously shown that it is possible to (compared to *in vitro* data-based *in silico* models) enhance the accuracy and coverage/range by approximately 3-fold with our software ANDROMEDA by Prosilico [1,5]). This software (available as SaaS and OnPrem; mainly for compounds with 150 to 750 g/mol molecular weight) is based on a large, unique human clinical PK-data base, machine learning and conformal prediction (CP) methodology. With this approach, valid confidence intervals are produced for each predicted numerical estimate.

This study was undertaken with the main aim to evaluate and compare the performance of 5 PK software (ANDROMEDA 2.0, and 4 free, web-based prediction tools; PkCSM, Swiss ADME, ADMETlab 3.0 and DruMAP 2.0) for predictions of f_a_ and f_u_ in humans. Compounds and data sets were selected in order to minimize the risk that test compounds have been used in training sets for model building (and thereby, influenced and exaggerated the predictive performances).

## Methods

The following software were used and compared for prediction of human f_a_ and f_u_ for sets of selected small compounds:

- **ANDROMEDA by Prosilico** 2.0 ([6] https://prosilico.com/andromeda/) – Only accessible by Prosilico and customers. Predicts f_a_ (numerical; considering passive permeability, efflux and dissolution), f_u_, steady-state volume of distribution (V_ss_), hepatic clearance (CL_H_), intrinsic hepatic metabolic CL (CL_int_), renal CL (CL_R_) biliary CL (CL_bile_) and total CL, MDR-1, BCRP, MRP2, oral bioavailability (F), half-life (t_½_), and many more parameters. Based on machine learning and conformal prediction (CP) methodology, which gives valid confidence in predictions (including confidence intervals for each predicted numerical estimate). Mainly applicable for compounds with MW 150 to 700 g/mol and for non-saturated conditions. Not applicable for metals and quaternary amines, and has limited use for hydrolysis sensitive compounds and covalently bound drugs.
- **PkCSM** ([7] https://biosig.lab.uq.edu.au/pkcsm/prediction) – Predicts f_a_ (numerical), f_u_, aqueous solubility, V_ss_, CL and MDR-1, but not parameters such as CL_H_, CL_R_, CL_bile_, F and t_½_.
- **SwissADME** ([8] http://www.swissadme.ch/) – Predicts S (insoluble to highly soluble), good or poor f_a_ (without defining how many % that is considered good) and MDR-1, but not parameters such as f_u_, V_ss_, CL_H_, CL_R_, CL_bile_, CL, F and t_½_.
- **ADMETlab** 3.0 ([9] https://admetlab3.scbdd.com/) – Predicts f_a_ (> or < 30 %), f_u_, V_ss_, *in vitro* CL_int_, MDR-1, CL and t_½_. but not but not parameters such as CL_H_, CL_R_ and CL_bile_.
- **DruMAP** 2.0 ([10] https://drumap.nibiohn.go.jp/) – Predicts f_a_ (low <20%; medium 20-70 %; high >70 %), f_u_, MDR-1, S, *in vitro* CL_int_, CL_R_ and f_e_-class, but not parameters such as CL_H_, CL_bile_, CL, V_ss_, F and t_½_.

The following data sets were selected and used:

a. A f_a_-data set with 44 compounds with varying permeability and dissolution (from very low to very high; including 9 compounds with f_a_ between ca 0 and 10 %, 16 compounds with f_a_<30 %, 15 compounds with limited dissolved fraction, and two apparent *in vivo* BCS IV-compounds), used for internal validation of ANDROMEDA by Prosilico (thus, not included in ANDROMEDA training sets) (Supplementary Table 1). Data were collected from the literature and internet.
b. A f_u_-data set with 54 compounds with very low f_u_ to 100 % f_u_, used for internal validation of ANDROMEDA by Prosilico (thus, not included in ANDROMEDA training sets) (Supplementary Table 2). Data were collected from the literature and internet.
c. A data set of 42 compounds (a majority not approved as medicines) for which human PK (apparently/probably) have not yet been disclosed and published (Supplementary Table 3; selected based on compounds highlighted by Drug Hunter). This reduces the risk of exaggerated prediction performance of the evaluated software.

Both quantitative (numerical) and qualitative (classes) evaluations and comparisons were done. For sets a and b, predicted results were compared to observed/measured estimates, which enabled evaluation of both actual and relative performances. For set c, for which measurements were unavailable, the relative performances of the software were evaluated.

The following classes and limits were selected:

f_a_ – low (<30 %; <0.3), moderate (30-80 %; 0.3-0.8) and high (>80 %:>0.8),

f_u_ – very low (<1 %; <0.01), low (1-3 %; 0.01-0.03), moderately low (3-10 %; 0.03-0.1), moderate (10-30 %; 0.1-0.3), moderately high (30-60 %; 0.3-0.6) and high (>60 %; >0.6).

In cases where values were below a certain level, 70 % of that was used for calculations of errors. For example, 0.007 was used for a compound with f_u_<0.01.

## Results

### Fraction absorbed (f_a_)

The f_a_-results are shown in Table 1, Figure 1 and Supplementary Tables 4 and 5.

**Table 1.** Prediction results for f_a_.

|  | ANDROMEDA | PkCSM | Swiss ADME | ADMETlab | DruMAP |
| --- | --- | --- | --- | --- | --- |
| *Prosilico test data set* |  |  |  |  |  |
| Q<sup>2</sup> | 0.69 | 0.35 | - | - | - |
| Mean absolute error (-fold) | 15 | 28 | - | - | - |
| Median absolute error (-fold) | 11 | 17 | - | - | - |
| Maximum absolute error (-fold) | 54 | 92 | - | - | - |
| Intercept (%; prediction axis) | 13 | 44 | - | - | - |
| Correct class prediction (%) | 77 | 45 | 23-34* | 16-84** | 64 |
| Large class prediction errors (%)*** | 0 | 9 | 8-24 | - | 0 |
| Failed predictions (%) | 0 | 2 | 11 | 0 | 2 |
| Inconclusive/uncertain (%) | 0 | 0 | 28 | 84**** | 0 |
| Software prediction overlap (%) | 9 |  |  |  |  |
| *Data set without available observed $f_a$-data* |  |  |  |  |  |
| Median $f_a$ (%) | 66 | 87 | - | - | - |
| Median $f_a$-class | Moderate | High | Low/Inconcl. | Mod/High | Moderate |
| Low $f_a$-predictions (%) | 2 | 2 | 38-71* | 0 | 14 |
| Software prediction overlap (%) | 24 |  |  |  |  |

\*Uncertain due to inconclusive predictions and prediction of two classes (good/poor); \*\*Uncertain because it predicts two classes (good/poor) only;
\*\*\*Predicted poor $f_a$ for high $f_a$ -compounds, or vice versa; \*\*\*\*Not possible to distinguish between moderate and high $f_a$ .

**Figure 1.**
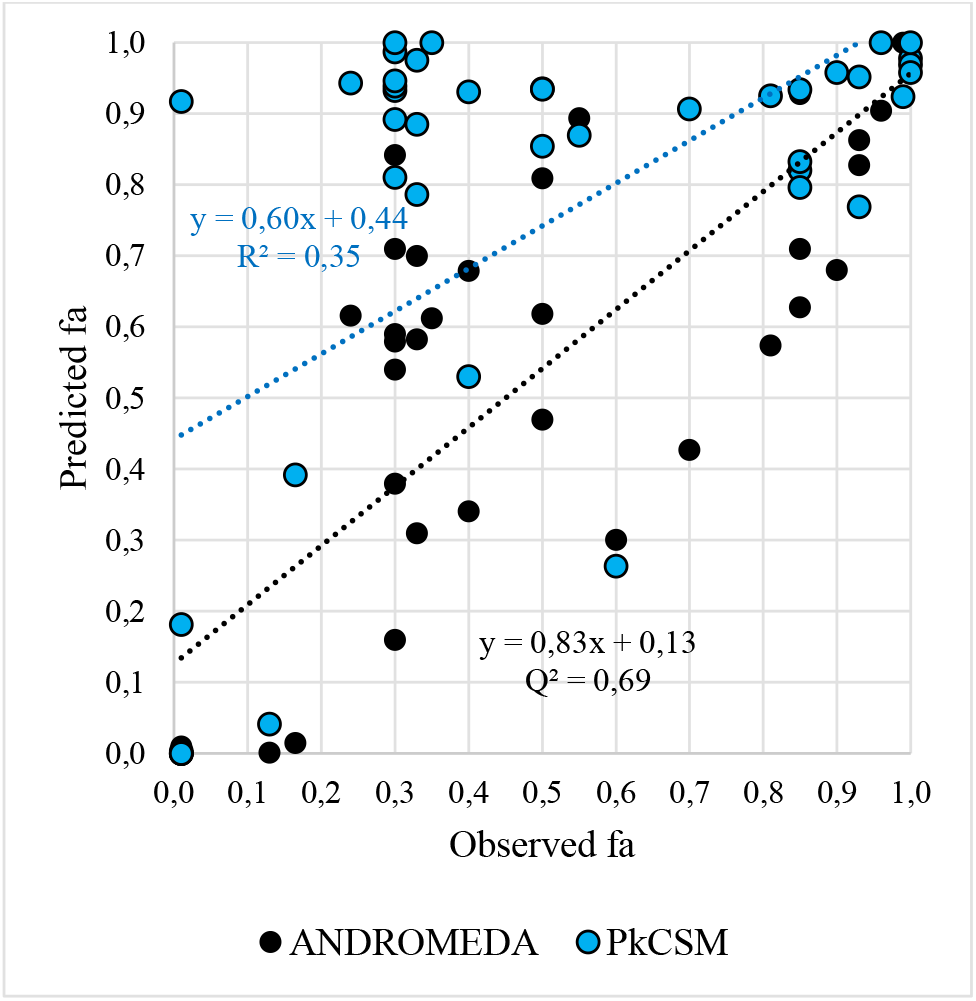
Observed f_a_ *vs* predicted f_a_ for the Prosilico test validation data set using ANDROMEDA and PkCSM (n=44).

#### Qualitative (numerical) predictions – Prosilico test validation data set

Q^2^-values (predicted *vs* observed f_a_) for ANDROMEDA and PkCSM were 0.69 and 0.35, respectively (Figure 1). Corresponding mean (median, maximum) absolute prediction errors were 15 (11, 54) % and 28 (17, 92) %, respectively. When using a default value of 50 % f_a_, mean, median and maximum predictions, absolute errors were 31, 34 and 50 %, respectively. Intercepts on the y-axis (predicted f_a_) were 13 and 44 %, respectively. For all 8 compounds with f_a_ close to zero (0-1 %), ANDROMEDA predicted zero f_a_, showing that it can predict poor absorption. PkCSM predicted zero uptake for 5 of them, but also 18 and 92 % f_a_ for two and failed prediction for one. For a compound with molecular weight below the domain for ANDROMEDA (AB13), f_a_ was underpredicted (1.5 *vs* 15-18 %).

#### Quantitative (classifications) predictions – Prosilico test validation data set

77 % correct predicted classes was reached with ANDROMEDA (low/moderate/high classes predicted). With PkCSM (low/moderate/high classes predicted) 45 % correct class predictions and 1 failure were found. Swiss ADME predicted classes of f_a_ and S, and for 32 % of compounds high f_a_ was predicted for insoluble or poorly soluble compounds. Thus, there was a high degree of inconsistency/uncertainty in these predictions. There were also 5 failed predictions with this software. At least 23 %, and maximally 34 %, of the class predictions were correct with Swiss ADME. ADMETlab predicts ><30 % f_a_, and 16-84 % (not possible to distinguish between moderate and high f_a_; can be approximated to average ca 50 %) of class predictions were correct. 3 of the 8 compounds with f_a_=0-1 % were predicted to have at least 30 % uptake. DruMAP predicted low f_a_ (<20 %) for these 8, but was unable to distinguish between zero and 19 % f_a_. With this software, 64 % of the predicted classes were correct and failed predictions were found for 1 compound.

For ANDROMEDA and DruMAP there were no predictions of low f_a_ for compounds with high f_a_, or vice versa. For PkCSM and Swiss ADME this occurred for 9 % and 8-24 % of predictions, respectively.

Overall, the 5 software predicted the same f_a_-class for 9 % of compounds only.

#### Quantitative (classifications) predictions – Data set without available observed f_a_-data

Median predicted f_a_ for ANDROMEDA and PkCSM were 66 (moderate) and 86 (high) %, respectively. Median predicted f_a_-class for Swiss ADME, ADMETlab and DruMAP were low/inconclusive, moderate/high (>30%) and moderate, respectively. For 24 % of compounds, the software predicted the same f_a_-class.

80 % of ANDROMEDA predictions were 35-78 % f_a_. Only one compound was predicted to have low f_a_ (28 %). PkCSM also predicted low f_a_ for one compound only (8 %). ADMETlab predicted >30 % f_a_ for all compounds. For DruMAP, 14 % of compounds were predicted to have low f_a_ (<20 %). Swiss ADME generated more low f_a_-predictions, 38 % (potentially up to 71%, when considering the 33 % inconsistent/uncertain predictions).

### Fraction unbound (f_u_)

The f_u_-results are shown in Table 2, Figures 2 and 3, and Supplementary Tables 6 and 7.

**Table 2.**
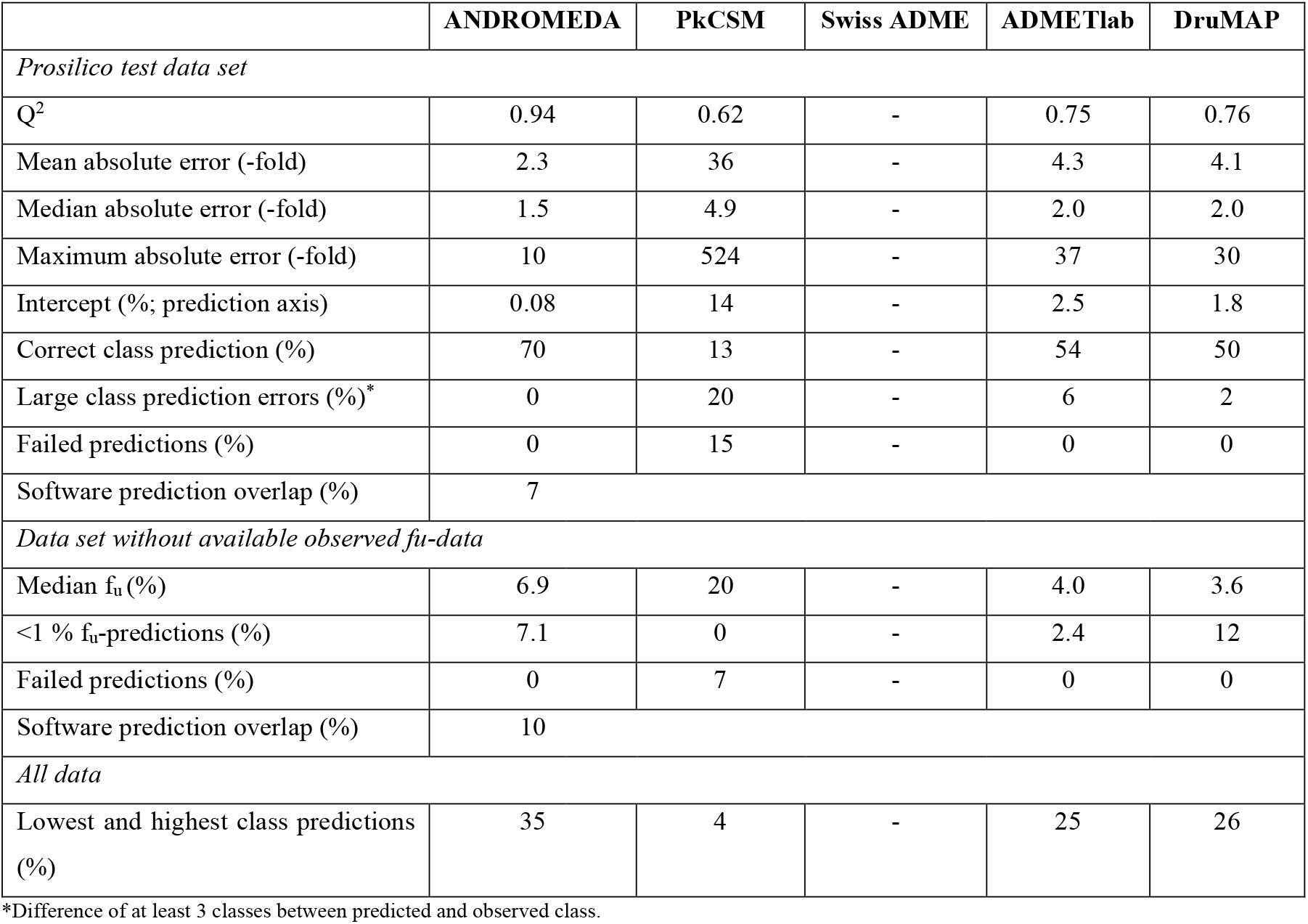
Prediction results for f_u_.

**Figure 2.**
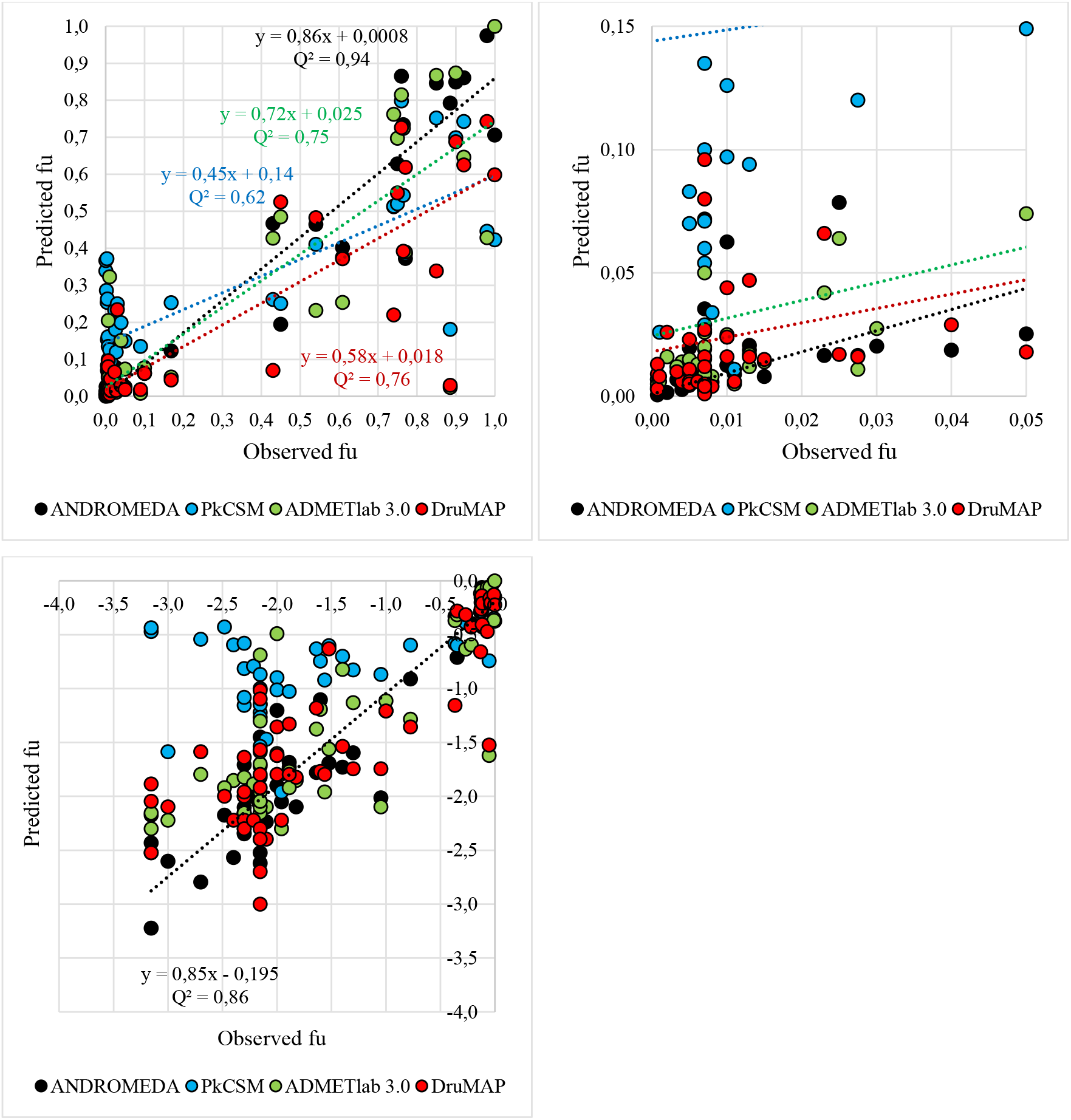
Observed f_u_ *vs* predicted f_u_ (left; full scale, right; 0.05-0.15 scale; below; logarithmic scale) for the Prosilico test validation data set using (n=54).

**Figure 3.**
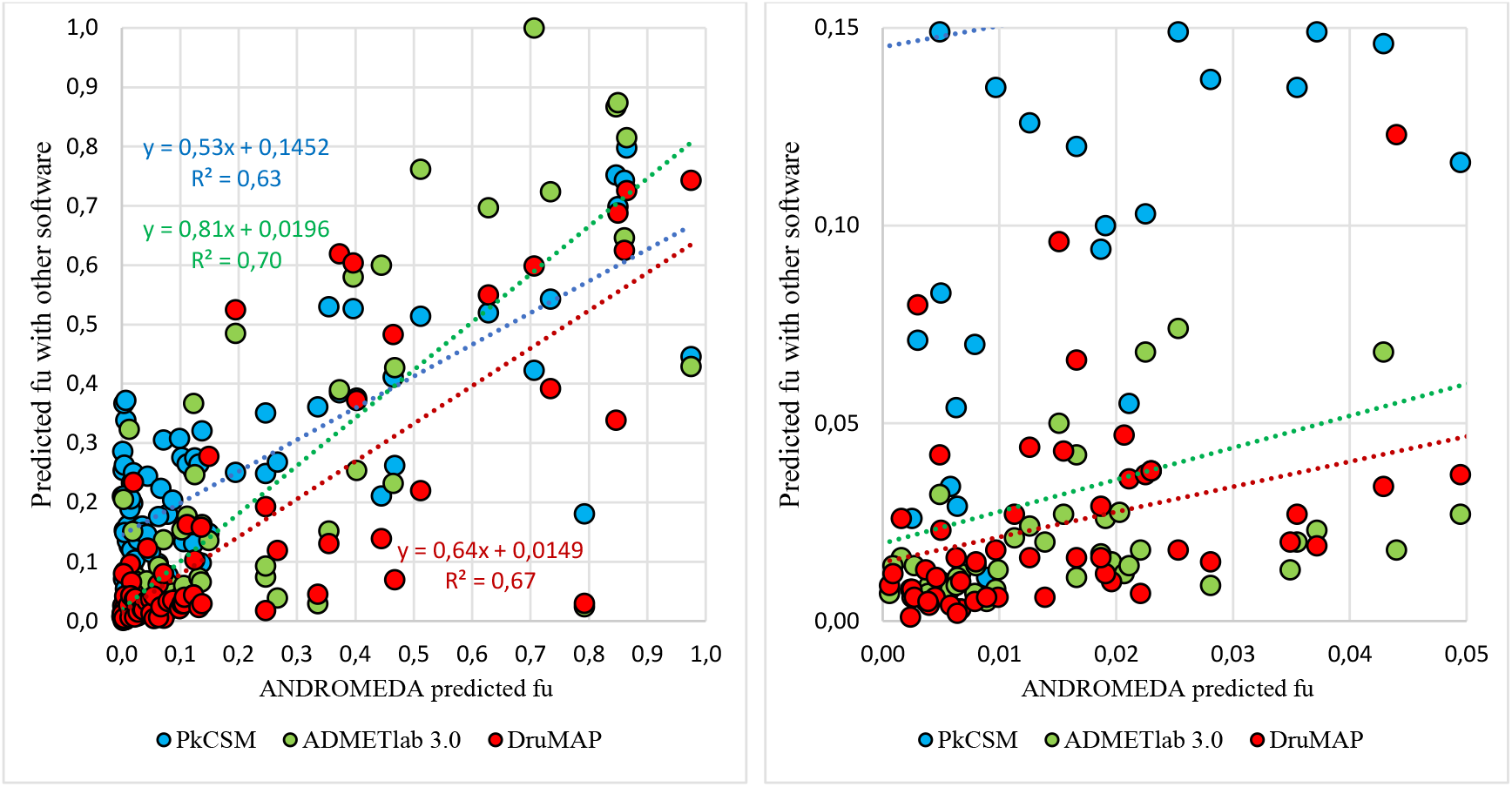
ANDROMEDA predicted f_u_ *vs* predicted f_u_ (left; full scale, right 0.05-0.15-scale) using PkCSM, ADMETlab 3.0 and DruMAP for all compounds (n=96).

#### Qualitative (numerical) predictions – Prosilico test validation data set

Q^2^-values (predicted *vs* observed f_u_) for ANDROMEDA, PkCSM, ADMETlab and DruMAP were 0.94, 0.62, 0.75 and 0.76, respectively (Figure 2). Corresponding mean (median, maximum) prediction errors were 2.3- (1.5-, 10-), 36- (4.9-, 524-), 4.3- (2.0-, 37-) and 4.1- (2.0-, 30-) fold, respectively. Intercepts on the y-axis (predicted f_u_) for the 4 software were 0.08, 14, 2.5 and 1.8 %, respectively.

R^2^-values for predictions with ANDROMEDA *vs* PkCSM, ADMETlab and DruMAP (all 96 compounds) were 0.63, 0.70 and 0.67, respectively (Figure 3). Corresponding intercepts on the y-axis (predicted f_u_) were 15, 2.0 and 1.5 %, respectively. PkCSM, ADMETlab and DruMAP also produced lower predictions than ANDROMEDA for compounds with higher f_u_, which was demonstrated by average predicted f_u_ of ca 65-80 % at 100 % with ANDROMEDA, respectively.

#### Quantitative (classifications) predictions – Prosilico test validation data set

Correct class was predicted for 70, 13, 54 and 48 % of compounds with ANDROMEDA, PkCSM, Swiss ADME and DruMAP, respectively. Corresponding numbers for large class prediction errors (≥3 classes difference between predicted and observed f_u_) were 0, 20 (and 15 % failed predictions), 6 and 2 %, respectively. The prediction overlap between software was 7 % (21 % for both data sets combined and when dividing f_u_ into 3 classes; <5 %, 5-50 % and >50 %).

#### Quantitative (classifications) predictions – Data set without available observed f_u_-data

For the data set without available observed f_u_-data there was 10 % prediction overlap between software. Median predicted f_u_ for ANDROMEDA, PkCSM, Swiss ADME and DruMAP were 6.9, 20, 4.0 and 3.6, respectively. PkCSM had 7 % failed predictions.

Overall (using both data sets), ANDROMEDA was the software with the highest percentage of predictions of lowest and highest f_u_-classes, 35 %, followed by DruMAP (26 %), ADMETlab (25 %) and PkCSM (4 %).

## Discussion

Five software were evaluated with regards to the ability to predict f_a_ and f_u_ for 140 test compounds in humans, and compared. Selected test compounds (including many with challenging absorption characteristics and very low f_u_) were not included in training sets for ANDROMEDA. It was not possible to explore whether this was also the case for the other software, but one can assumed that at least parts of selected test compounds have been used in training sets for these. In part, test compounds were selected in order to minimize this risk.

For both f_a_ and f_u_, **ANDROMEDA** was clearly more accurate than the other investigated software, with higher Q^2^ (0.69 vs 0.35 and 0.94 *vs* 0.62-0.76), lower mean errors (15 % *vs* 28 % and 2.3- *vs* 4.1- to 36-fold), lower maximum errors (54 % *vs* 92 % and 10- *vs* 30- to 524-fold), more correct predicted classes (70-77 % *vs* 13-54 %), no failed, inconclusive or poor predictions (as found for 3 of the other software), wider application range, and minimal skewness at low values. It had intercepts on f_a_- and f_u_-prediction axes that were ca 1/3 and 1/175 to 1/23 compared to those found for the other software. It correctly predicted the f_a_ of the 8 compounds with poorest absorption, and shows 1 % mean absolute prediction error for all new compounds with 0-5 % f_a_ added to the upcoming software version (ANDROMEDA 3.0). The performance was in agreement with previous studies. For example, corresponding Q^2^, mean error and intercept obtained in a study with 136 compounds, including 59 that violate the Rule of 5, were 0.72, 14 % and 13 %, respectively [3]. Somewhat higher f_a_ (ca 0.8) and lower intercept (6 %) have been found in other validation studies with ANDROMEDA.

ANDROMEDA 3.0 has 6 % better performance (higher Q^2^, lower mean error and more narrow confidence intervals) than version 2.0 for the prediction of both f_a_ and f_u_ in humans, and has ca ¼ lower f_u_-intercept. which further increases the lead. Another advantage with this software is the production of true confidence intervals for each compound and parameter estimate.

Advantages with ANDROMEDA suggest that this is the software of choice for those that desire adequate predictions of f_a_ and f_u_ in humans and estimates of certainty.

Reasons to the better performance of ANDROMEDA are believed to include its validated prediction models for passive permeability-based uptake, efflux and fraction dissolved, numerical scale, unique algorithms and large amount of biopharmaceutical data used for model building.

Poorest overall performance was found for **PkCSM**, with large overpredictions (up to 524-fold; also for its V_ss_-model) and many failed predictions. It predicted near complete to complete absorption for 11 compounds with 0 to 30 % uptake, and reached a mean prediction error for f_a_ that was of same magnitude as that obtained when assuming 50 % f_a_ for all compounds. Thus, with this software there is high risk to overpredict absorption and CL, be misled in the predictions of bioavailability and t_½_, and make inadequate decisions.

**Swiss ADME** was associated with many failed and inconclusive/contradictory predictions for f_a_, and was not applicable for f_u_-predictions. It produced even weaker f_a_-predictions than PkCSM, with high portions of inconclusive, low and failed predictions, and the lowest percentage of correct class predictions, only 23-34 %. A significant proportion of compounds were predicted to have both high f_a_ and poor solubility or insolubility, and this creates confusion and uncertainty. Two compounds were predicted to be insoluble with Swiss ADME, but to have high f_a_ according to predictions with all 5 software and confirmed high *in vivo* f_a_. It was, however, relatively accurate in predicting low f_a_ for 10 of 18 poorly absorbed compounds. For the remaining 8 it predicted high f_a_. The results speak against Swiss ADME as an accurate, consistent and trustworthy software for prediction of f_a_ in humans.

**ADMETlab** took a middle position performance-wise, and is limited by its 2.5 % f_u_-intercept (overprediction potential of highly bound compounds; see below), case with poor prediction (37-fold underprediction of f_u_ for one compound), and dichotomous f_a_-model (> or < 30 %; without ability to distinguish between very poor and moderately low f_a_ and between moderately high to very high f_a_). A major limitation with ADMETlab is the extensive underprediction trend for t_½_, with predicted t_½_ ranging from ca 0.2 to ca 2 h (a majority <1 h) for a large set of compounds with adequate t_½_ in humans.

Apparently, **DruMAP** was second best performing software. Drawbacks with this software include the 3-model for f_a_ (low, moderate & high; without ability to distinguish between very poor and moderately low f_a_ and between moderately high to very high f_a_) and cases with poor f_a_- and f_u_- predictions (up to 30-fold error). It is not known whether this was due to comparably large number of training data set compounds among test compounds.

**ADMET Predictor** was not included in this evaluation since it is not available as a freeware on the web. It has previously been shown to have an intercept of ca 6 % for f_u_-predictions (implying risk of extensive overpredictions for highly bound compounds) (Yun). Published f_a_-prediction results for ADMET Predictor are limited. The intermediate parameter used for ADMET Predictor predictions of absorption, effective human intestinal permeability (P_eff_), has been shown to have a LOQ in the moderately high permeability zone (corresponding to ca 70-90 % f_a_), which is a considerable limitation [1].

**Intercepts** on prediction axes are expected with *in silico* models, and the greater they are, the more impact they will have on the predictions (of for example, CL, t_½_ and F) and decision-making. Many modern small drugs have low f_u_ and limited f_a_ [5], implying that skewed models are likely to mislead siginficantly. For a compound with 1 % f_u_, the general overprediction trend is 1.3-, 40-, 2.6- and 20-fold (estimated using logged values) with ANDROMEDA, PkCSM, ADMETlab and DruMAP, respectively. Corresponding estimates for a compound with a f_u_ of 0.1 % are 1.9-. 394-, 7.0- and 194-fold, respectively.

Overall, there was only 7-24 % **performance overlap** between the 5 software. For example, 60 % of investigated compounds were predicted to have both poor and good f_a_. There were cases where one software predicted very low f_u_ and another moderate f_u_ (with 233-fold difference) and a case where low and high f_u_ was predicted with different software (2.4 and 79 %). This will, of course, be confusing for those who use all these for predictions of the selected parameters. The discrepancy could partly be explained by inclusion of many test compounds with complex absorption and low f_a_ and very low f_u_ (=difficult to predict), and partly by poor performances among tested software. With the use of only one validated software with adequate performance (such as ANDROMEDA) such a problem could be removed/reduced.

For those interested in using free or purchased tools or internally developed software, and concerned about clinical accuracy, validity, trustworthiness and application domains, it is recommended to request performance results, including a broad range of statistics obtained in cross-validations and hold-out set tests relevant for humans *in vivo*. Software developers concerned about proprietary rights (as we are) are unlikely to reveal their training data sets and algorithms, and this is a major issue for external investigations, including this one. In this study we selected test sets that we have not used for model building, which enabled us to present true own predictions. Prediction results for the 42 compounds for which PK are not yet available (at to our knowledge) can easily be checked by developers of them and others.

## Conclusion

A handful of software were evaluated with regards to the ability to predict human f_a_ and f_u_ for 140 test compounds with varying characteristics, and also compared. Test compounds were not included in training sets for ANDROMEDA. It was, however, not possible to explore whether this was also the case for the other software. For both f_a_ and f_u_, ANDROMEDA was clearly more accurate and balanced than the others. The higher performance, in particular, in the lower ranges of f_a_ and f_u_, are of great value and importance. Two software, PkCSM and Swiss ADME, were considered inappropriate, whereas ADMETlab took an intermediate performance position. Apparently, DruMAP was second best performing software. Overall, there was poor performance overlap between the software, with many contradictory predictions. Challenging PK-properties and software with poor performance are among explanations. Advantages with ANDROMEDA suggest that this is the software of choice for those that desire adequate predictions of f_a_ and f_u_ in humans.

---

**Suppmentary Table 1.** 44 compounds from Prosilico’s internal validation data set for f_a_.

|  |  | fa in man |
| --- | --- | --- |
| AB1 | CCc1cc2c(cc1N3CCC(CC3)N4CCOCC4)C(c5c(c6ccc(cc6[nH]5)C#N)C2=O)(C)C | 0,5 |
| AB2 | C[C@]3([H])O[C@@]([H])O[C@@]1([H])[C@](O)([H])[C@@]([H])O[C@]1([H])CO)O[C@@]2([H])[C@](O)([H])[C@@]([H])O[C@]2([H])CO)O[C@]1([H])O[C@@]([H])O[C@]3([H])N[C@]4([H])C=C(CO)[C@@]([H])O[C@@]([H])O[C@@]4([H]) | 0,01 |
| AB3 | NCC[C@](O)([H])C(=O)N[C@]2([H])C[C@@]([H])([H])[C@]1([H])O[C@@]([H])(CN)[C@](O)([H])[C@@]([H])[C@@]1(O)[H])[C@@]([H])[C@]2([H])O[C@]3([H])O[C@@]([H])(CO)[C@](O)([H])[C@@]([H])[C@@]3(O)[H] | ca 0 |
| AB4 | CC[C@H]1OC(=O)[C@H](C)[C@@H](O[C@H]2C[C@@](C)(OC)[C@@H](O)[C@H](C)O2)[C@H](C)[C@@H](O)[C@@H]2O[C@H](C)C[C@H]([C@H]2O)N(C)C)[C@](C)(O)C[C@@H](C)CN(C)[C@H](C)[C@@H](O)[C@]1(C)O | 0,60 |
| AB5 | CCS(=O)(=O)N1CC(CC#N)(n2cc(-c3ncnc4[nH]ccc34)cn2)C1 | 0,85 |
| AB6 | CC1=C(C(CCC1)(C)C)C=CC(=CC=CC(=CC=CC(C)C)C=CC(C)C)C=CC2=C(CCCC2(C)C)C)C | ca 0 |
| AB7 | CC(C)C(C)C1(CCC2(C3CCC4C5(COCC4(C3=CCC2(C1C(=O)O)C)CC(C5OCC(C)(C(C)(C)C)N)N6C(=NC=N6)C7=CC=NC=C7)C)C)C | 0,70 |
| AB8 | NCCCC(O)(P(=O)(O)O)P(=O)(O)O | ca 0 (very low Papp) |
| AB9 | Cn1c2c(c3ccccc31)C(=O)C(Cn1ccnc1)CC2 | ca 1 (high Papp) |
| AB10 | COC(=O)C1=C(C)NC(C)=C(C(=O)OC(C)C)C1c1cccc2nnc12 | ca 1 (high Papp) |
| AB11 | CS(=O)(=O)Nc1cc2c(c(oc2cc1)CCCC)C(=O)c1ccc(cc1)OCCCN(CCCC)CCCC | 0,33 |
| AB12 | CC[C@@H]1[C@@]([C@@H]([C@H](C(=O)[C@@H](C[C@@]([C@@H]([C@H]([C@@H]([C@H](C(=O)O)1)C)O[C@H]2C[C@@]([C@H]([C@@H](O2)C)O)(C)OC)C)O[C@H]3[C@@H]([C@H]([C@H](O3)C)N(C)C)O)(C)O)C)O)(C)O | 0,40 |
| AB13 | O=C(O)P(=O)(O)O | 0,15-0,18 |
| AB14 | [C@@]67(O)[C@H]1[C@@H]([C@@]5([C@](O)(CC1)C[C@@H](O[C@@H]4O[C@@H]([C@@H](O[C@@H]3O[C@@H]([C@@H](O[C@@H]2O[C@@H]([C@@H](O)[C@H](O)[C@H]2O)CO)[C@H](O)[C@H]3O)CO)[C@@H](OC)C4)CC5)C=O)CC[C@@]6([C@H](CC7)C8=CC(OC8)=O)C | 0,10-0,16 |
| AB15 | C(C1C(C(C(C(O)OC(C(CO)O)C(C(CO)O)O)O)O)O | ca 0 |
| AB16 | CCOC(=O)N1CCC(=C2C3=C(CCC4=C2N=CC=C4)C=C(C=C3)Cl)CC1 | 0,90 |
| AB18 | N[C@@H]1c2ccnnc2[C@H](OC(=O)N2CCC(n3c(=O)[nH]c4ncccc43)CC2)CC[C@H]1c1cccc(F)c1F | 0,81 |
| AB19 | CC(C)C[C@@H]1CN2CCc3cc(c(cc3[C@H]2C[C@H]1OC(=O)[C@H](C(C)C)N)OC)OC | 0,55 |
| AB20 | CC1CC2C3CCC4=CC(=O)C=CC4(C3=CCC2(C1(C(=O)CO)O)C)C | ca 1 |
| AB21 | C[C@H](COC1=CC=C(C=C1)C2=CN=C3N2N=C(C=C3)N[C@H](C)C4=CC(=CC=C4)F)N | >0,7-0,8 |
| AB22 | CCCCN(CCCC)C(=O)CN1C[C@@H]([C@H]([C@@H]1C1=CC=C(OC)C=C1)C(O)=O)C1=CC2=C(OCO2)C=C1 | >0,94 |
| AB23 | CC1=C(C(=O)OC2=C1C=CC(=C2)OC3=NC=CC=N3)CC4=C(C(=NC=C4)NS(=O)(=O)NC)F | >0,9 |
| AB24 | CS(=O)(=O)CCNCC1=CC=C(O1)C2=CC3=C(C=C2)N=CN=C3NC4=CC(=C(C=C4)OCC5=CC(=CC=C5)F)C1 | 0,35 |
| AB25 | CC1(CCC2(CCC3(C(C2C1)C(=O)C=C4C3(CCC5C4(C=C(C(=O)C5(C)C)C#N)C)C)NC(=O)C(C)(F)F)C | 0,23-0,25 |
| AB30 | CCCS(=O)(=O)N[C@H]1C[C@H](C1)N(C)C1=C2C=CNC2=NC=N1 | ca 0,5 |
| AB31 | CC1(CCC2C(C1)CCC3C2CCC4(C3CCC4C(=O)CN5C=C(C=N5)C#N)C)O | ca 1/3 |
| AB32 | [H][C@]12SCC(C[N+]3(CCNC(=O)C4=C(Cl)C(O)=C(O)C=C4)CCCC3)=C(N1C(=O)[C@H]2NC(=O)C(=N/OC(C)(C)C(O)=O)C1=CSC(N)=N1)C([O-])=O | Very low (close to 0) |
| AB33 | ClCCC(c1ccc(cc1)CCCC(=O)O)CCCCl | 0,85 |
| AB34 | C1CN(CCN1CCC=C2C3=CC=CC=C3SC4=C2C=C(C=C4)Cl)CCO | 0,93 |
| AB35 | CCC(C)CCCC(=O)NC(CCN)C(=O)NC(C(C)O)C(=O)NC(CCN)C(=O)NC1CCNC(=O)C(NC(=O)C(NC(=O)C(NC(=O)C(NC(=O)C(NC(=O)C(NC1=O)CCN)CC(C)C)CC(C)C)CCN)CC(C)O | Very low (close to 0) |
| AB36 | O1[C@@H](C)[C@H](O)[C@@H](O)[C@@H](O)[C@@H]1O[C@H]1[C@H](OC(=O)C=Cc2cc(O)c(O)cc2)[C@H](O)[C@@H](OCCc2cc(O)c(O)cc2)[C@@H]1O)CO[C@@H]1O[C@H](CO)[C@@H](O)[C@H](O)[C@H]1O | Very low (close to 0) |
| AB37 | C[C@H]1CN(C(=O)CC#N)[C@]11CCN(C1)C1=C2C=CNC2=NC=N1 | >0,7-0,8 |
| AB38 | CCCCC1=CC(=C2[C@H]3C=C(C(C[C@H]3C(OC2=C1)(C)C)C)O | ca 0,4 |
| AB39 | CC1C=CC=C(C(=O)NC2=C(C3=C(C4=C(C(=C3O)C)OC(C4=O)(OC=CC(C(C(C(C(C1O)C)O)C)OC(=O)C)C)OC)C)C5=C2N6C=CC(=CC6=N5)C)O)C | ca 1/3 |
| AB40 | OC(=O)C=C(C=C(C=C(C=C/C=C1C(CCCC=1C)(C)C)C)/C | 0,50 |
| AB41 | C4=C(C1CCC(CC1)C3=C(O)C2=CC=CC=C2C(=O)C3=O)C=CC(=C4)Cl | <0,3 |
| AB42 | CCCSC1=CC2=C(C=C1)N=C(N2)NC(=O)OC | <0,3 |
| AB44 | CCCS(=O)(=O)Nc1ccc(F)c(c1F)C(=O)c2c[nH]c3c2cc(cn3)c4ccc(Cl)cc4 | 0,30 |
| AB45 | CC1(C)C[C@H]2CN1C1=NC(=CC=C1C(=O)NS(=O)(=O)C1=CC=CC(NCCC2)=N1)N1C=CC(OC2C3(C(C3)C2CC2)=N1 | 0,3 |
| AB46 | [2H]C([2H])([2H])C(C1=CC(=C(NC(=O)C2=CNC3=C(C=CC=C3)C2=O)C=C1O)C(C)(C)C(C([2H])([2H])[2H])C([2H])([2H])[2H] | 0,3 |
| AB47 | CC1=C(C(CCC1)(C)C)C=CC(=CC=CC(=CC(=O)NC2=CC=C(C=C2)O)C)C | ca 0,3 |
| AB48 | CC[C@H](C(=O)OC1CC(C)C=C2[C@H]1[C@@@H](CC[C@@H]1C[C@@@H](O)CC(=O)O1)[C@H](C=C2)C)C | ca 0,3 |
| AB49 | CCOC(=O)C1=C(NC(=C(C1C2=C(C(=CC=C2)Cl)Cl)C(=O)OC)C)C | Near complete |
| AB50 | CC1CC(=O)C=C(C12C(=O)C3=C(O2)C(=C(C=C3OC)OC)Cl)OC | ca 1 |

**Suppmentary Table 2.** 54 compounds from Prosilico’s internal validation data set for f_u_.

|  |  | f<sub>u</sub> |
| --- | --- | --- |
| D1 | CC1=C(C(=CC=C1)F)C(=O)N2CCCC(C2C3=CC=C(C=C3)NC4CCCC4)C(=O)NC5=CC(=C(C=C5)C)C(F)(F)F | <0,001 |
| D2 | COc1cc(C)c(C=CC(C)=CC=CC(C)=CC(O)=O)c(C)c1C | 0,005 |
| D3 | O=C1NC(=O)CCC1(c2ccc(N)cc2)CC | 0,77 |
| D4 | O=C3OCC(O)C3=CCC1\C(=C)CCC2C(C)(CO)C(O)CCC12C | 0,45 |
| D5 | NCC1=CC(=CC=C1)N1N=C(C=C1C(=O)NC1=CC(=CC=C1F)[C@H](NCC1CC1)C1=CC=CC(=C1)C#N)C(F)(F)F | 0,01 |
| D6 | NC(C(=O)NC1C2SCC(Cl)=C(N2C1=O)C(O)=O)c1cccc1 | 0,75 |
| D7 | CO/N=C(/C1=NSC(=N1)N)\C(=O)N[C@H]2[C@@H]3N(C2=O)C(=C(CS3)C[N+](45CCCC(CC4)(CC5)C(=O)N)C(=O)[O-]) | 0,92 |
| D8 | C1CC1C(=O)NC2=NN3C(=N2)C=CC=C3C4=CC=C(C=C4)CN5CCS(=O)(=O)CC5 | 0,43 |
| D9 | CC(=O)[C@H]1CC[C@H]2[C@@]1(CC[C@H]3[C@H]2CC[C@H]4[C@@]3(CC[C@@](C4)(C)O)C)C | ca 0,01 |
| D10 | COc1ccc(Cl)cc1C(=O)NCCc1ccc(cc1)S(=O)(=O)NC(=O)NC1CCCCC1 | 0,001 |
| D11 | CC(O)C(C)N1N=CN(C1=O)C1=CC=C(C=C1)N1CCN(CC1)C1=CC=C(OC2COC(CN3C=NC=N3)(O2)C2=C(C)C=C(Cl)C=C2)C=C1 | 0,005 |
| D12 | C[C@H](C(C)C)[C@]1(CC[C@@]2([C@H]3CC[C@H]4[C@]5(COC[C@]4(C3=CC[C@]2([C@@H]1C(=O)O)C)C[C@H]([C@@H]5OC[C@@](C)(C(C)C)N)N6C(=NC=N6)C7=CC=NC(=C7)C)C)C | <0,01 |
| D13 | CS(=O)(=O)CCNCC1=CC=C(O1)C2=CC3=C(C=C2)N=CN=C3NC4=CC(=C(C=C4)OCC5=CC(=CC=C5)F)Cl | 0,005 |
| D14 | Clc1ccc(cc1)\C=C3c4c(c2c(cc(Cl)cc23)C(O)CN(CCCC)CCC)ccc(Cl)c4 | <0,01 |
| D15 | CC1=C(C)C(NC2=C(C=CC=C2)C(O)=O)=CC=C1 | 0,05 |
| D16 | CN1C=CN1=S | 0,90 |
| D17 | CC1=C(C(=O)C2=C(C1=O)N3C[C@H]4[C@@H]([C@@]3([C@H]2COC(=O)N)OC)N4)N | 0,76 |
| D18 | ClC1=CC(Cl)=C(C=C1)[C@]3(CN2C=NC=N2)OC[C@H](COC4=CC=C(C=C4)N5CCN(CC5)C6=CC=C(C=C6)N7C=NNC7=O)O3 | 0,011 |
| D19 | CN1CCC23C4OC5=C2C(CC1C3C=CC4O)=CC=C5OCCN1CCOCC1 | 0,77 |
| D20 | CCc1ccc(CCOc2ccc(CC3SC(=O)NC3=O)cc2)nc1 | 0,005 |
| D21 | CC(=O)O[C@H]1C[C@H]2CC[C@H]3[C@H]([C@]2[C@H]1N4CC[N+](CC4)(C)C)CC[C@]5([C@H]3C[C@H]([C@@H]5OC(=O)C)N6CC[N+](CC6)(C)C)C | 0,98 |
| D23 | CC(C)c4nc(CN(C)C(=O)N[C@H](C(C)C)C(=O)N[C@H](Cc1ccccc1)C[C@H](O)[C@H](Cc2ccccc2)NC(=O)OCc3cncs3)cs4 | 0,015 |
| D24 | CN1C(=O)N2C=NC(=C2N=N1)C(=O)N | 0,85 |
| D25 | O=C(Nc1noc(c1)C)Nc4ccc(Oc2c3cc(OC)c(OC)cc3ncc2)cc4C1 | <0,01 |
| D26 | CC(C)OC1=C(C=C(C=C1)C2=NN(C3=NC=NC(=C23)N)C(C)C4=C(C(=O)C5=C(O4)C=CC(=C5)F)C6=CC(=CC=C6)F)F | <0,003 |
| D27 | CCCCC(=O)OCC(=O)[C@]1(C[C@H](C2=C(C1)C(=C3C(=C2O)C(=O)C4=C(C3=O)C=CC=C4OC)O)O[C@H]5C[C@H]([C@H]([C@@H](O5)C)O)NC(=O)C(F)(F)F)O | 0,005 |
| D28 | Cc1ccccc1S(=O)(=O)NC(=O)c2cc(OC)c(cc2)Cc3cn(C)c4ccc(cc43)NC(=O)OC5CCCC5 | <0,01 |
| D37 | C[C@H]1CN(C(=O)CC#N)[C@]11CCN(C1)C1=C2C=CNC2=NC=N1 | 0,71-0,78 |
| D38 | C[C@H](COC1=CC=C(C=C1)C2=CN=C3N2N=C(C=C3)N[C@H](C)C4=CC(=CC=C4)F)N | 0,04 |
| D39 | CCCCN(CCCC)C(=O)CN1C[C@@H]([C@H]([C@@H]1C1=CC=C(OC)C=C1)C(O)=O)C1=CC2=C(OC(O2)C=C1 | <0,01 |
| D40 | CNC(=O)C1=CC=C(C=C1)NC2=NC=C(C(=N2)NCC3=NC=CN=C3N(C)S(=O)(=O)C)C(F)(F)F | 0,1 |
| D41 | CC1=C(C(=O)OC2=C1C=CC(=C2)OC3=NC=CC=N3)CC4=C(C(=NC=C4)NS(=O)(=O)NC)F | 0,01 |
| D42 | CO[C@H]1\C=C\O[C@]2(C)Oc3c(C)c(O)c4c(O)c(NC(=O)\C=C/C=C/[C@H](C)[C@H](O)[C@H](C)[C@H](O)[C@H](C)[C@H](OC(=O)C)[C@@H]1)C)c(\C=N\N5CCN(CC5)C6CCCC6)c(O)c4c3C2=O | 0,023 |
| D43 | C[C@H]1O[C@H](C[C@H](O)[C@@H]1O)O[C@H]2[C@@H](O)C[C@H](O[C@H]3[C@H](O)C[C@H](O[C@H]4CC[C@H]5C)[C@H](CC[C@@H]6[C@@H]5CC[C@]7(C)[C@H](CC[C@]67O)C8=C(C(=O)OC8)C4)O[C@@H]3CO[C@H]2C | 0,030 |
| D44 | CCCSe1ccc2[nH]c(NC(=O)OC)nc2c1 | 0,169 |
| D45 | OP(=O)(O)OC(Cn1cncn1)(Cn2cncn2)c3ccc(F)cc3F | 0,885 |
| D46 | Cc1cccc(C)c1NC(=O)CC23CCCN2CCC3 | 0,540 |
| D47 | CCN(CC)CCNC(=O)c1cc(ccc1OC)S(=O)(=O)C | 1,000 |
| D48 | CCCN(CCC)C(=O)CC1=C(N=C2N1C=C(C=C2)Cl)C3=CC=C(C=C3)Cl | 0,0061 |
| D49 | NC(=O)OC[C@H](O)C1=CC=CC=C1Cl | 0,609 |
| D50 | OC(=O)C1=CC=NC(=C1)N1CC(O)(C1)C1=C(Cl)C=C(OCC2=C(ON=C2C2=C(Cl)C=CC=C2Cl)C2CC2)C=C1 | 0,004 |
| D51 | [H][C@@]12CN(C(=O)[C@@]([H])(NC(=O)O[C@@]3([H])CCC[C@@]3([H])OC\C=C\C(F)(F)C3=NC4=C(C=CC=C4)N=C3O1)C(C)(C)C)[C@@]([H])(C2)C(=O)N[C@@]1(C[C@H]1C(F)F)C(=O)NS(=O)(=O)C1(C)CC1 | 0,0275 |
| D52 | OC(=O)C[C@H]1CC[C@@H](CC1)C1=CC=C(C=C1)C1=NC=C(NC2=CN=C(C=C2)C(F)(F)F)C=C1 | 0,008 |
| D66 | CCCCCCCCC[C@H](C[C@@H]1OC(=O)[C@H]1CCCCC)OC(=O)[C@H](CC(C)C)NC=O | <0,01 |
| D68 | ClC1=CC2=C(C=C1)C(=CC=N2)N1CCN(CCCN2CCN(CC2)C2=CC=NC3=C2C=CC(Cl)=C3)CC1 | <0,01 |
| D69 | N([C@H]([C@H](C[C@H](Cc1ccccc1)NC(=O)[C@@H](N1C(=O)NCCCC1)C(C)C)O)Cc1ccccc1)C(=O)COc1c(cccc1C)C | <0,02 |
| D70 | CC1=C(C=CC=C1O)C(=O)NC(CSC2=CC=CC=C2)C(CN3CC4CCCCC4CC3C(=O)NC(C)(C)C)O | <0,02 |
| D71 | CC#CC(=O)N1CCC[C@H]1C1=NC(=C2N1C=CN=C2N)C1=CC=C(C=C1)C(=O)NC1=CC=CC=N1 | 0,025 |
| D72 | COC1=NC2=C(C=C(Br)C=C2)C=C1[C@@H](C1=CC=CC=C1)[C@@](O)(CCN(C)C)C1=CC=CC2=C1C=CC=C2 | <0,001 |
| D73 | [H][C@]1(CCCN1C(=O)[C@@H](NC(=O)OC)C(C)C)C1=NC=C(N1)C1=CC2=C(C=C1)N1[C@@H](OC3=C(C=CC(=C3)C3=CN=C(N3)[C@]3([H])CCCN3C(=O)[C@@H](NC(=O)OC)C(C)C)C1=C2)C1=CC=CC=C1 | <0,001 |
| D74 | COC1=CC2=NC3=C(CCCCC[C@@H]4C[C@H]4OC(=O)N[C@H](C(=O)N4C[C@@H](C[C@H]4C(=O)N[C@@]4(C[C@H]4C=C)C(=O)NS(=O)(=O)C4CC4)O3)C(C)(C)C)N=C2C=C1 | <0,012 |
| D75 | ClC1=CC(Cl)=C(C=C1)[C@@H]1CS\C(S1)=C(\C#N)N1C=CN=C1 | <0,01 |
| D76 | [H][C@@]1(CC[C@@]2([H])[C@]3([H])[C@H](O)[C@H](CC)[C@]4([H])C[C@H](O)CC[C@]4(C)[C@@]3([H])CC[C@]12C)[C@H](C)CCC(O)=O | <0,01 |
| D78 | COC[C@H]1C[C@H](N(C1)C(=O)[C@H](NC(=O)OC)C1=CC=CC=C1)C1=NC=C(N1)C1=CC=C2C(COC3=CC4=C(C=CC5=C4NC(=N5)[C@@H]4CC[C@H](C)N4C(=O)[C@@H](NC(=O)OC)C(C)C)C=C23)=C1 | <0,005 |

**Suppmentary Table 3.** 42 compounds without published f_a_ and f_u_-values.

|  |  |
| --- | --- |
| ALG000184 | O=C(NC1=CC=C(F)C(C#N)=C1)C2=C(C)C(C(C(N(COP(O)(O)=O)C3(COC3)C#C)=O)=O)=C(C)N2C |
| Alixorexton | [H][C@]1(C2=CC=CC=C2OCC3=O)CC[C@@]([H])(OC[C@H]4[C@@H](NS(C)(=O)=O)CCCN34)CC1 |
| AMG410 | FC1=C(C2=C(C=NN3)C3=CC(C1)=C2CCCO4)N=CC5=C1N=C(OC[C@]67C[C@@H](F)CN6CCC7)N=C5N8CCOC[C@@H](OC4=O)C8 |
| ARN23746 | O=C(O)C1=CC=C(NCCCCCCCC(F)(F)F)C(S(=O)(N(C)C)=O)=C1 |
| AZD0022 | FC1=C(C#C)C2=[C@]([C@]3=C(F)C(N=C(OC[C@]4[C[C@@H](F)C5)N5CCC4)N=C6N7C[C@H]8N[C@H](C=C8)C7)=C6C=C3F)C=C(N)C=C2C=C1 |
| AZD0780 | O=C1C=CC=CN1C(C=N2)=CC=C2N[C@H](C3)CC[C@@H]3NC4=NC=C(C=N4)OC(F)F |
| AZD2389 | O=C(N1[C@@H](CSC1)C#N)CNC(C2=CC=NC3=C2C=C(N4CCOCC4)C=C3)=O |
| Azetukalner | Cc1cc(N2CCc3cc(F)ccc3C2)cc(C)c1NC(=O)CC(C)(C)C |
| BAY3389934 | CN1C=CN=C1CCOC([C@@H](NS(C2=C(CC)C(N3C([C@@H](O)CC3)=O)=CC=C2)(=O)=O)CNC(C4=CC=C(C1)S4)=O)=O |
| BBO10203 | O=C(C=C)N1[C@H](C)C2=CC(C3=[C@]([C@]4=C(F)C=C(F)C=C4OCCOC)C(SC=C5)=C5C(C6=CC=C(N(C)N=C7)C7=C6)=N3)=NN2CC1 |
| BI9508 | CC(C)OC1=CC=C(C2=C3C(N(C)N=C3)=C(N(CCOC(F)F)C([C@H]4[C@@H](C5=CC=NC=N5)C4)=O)C=C2)C=C1 |
| BMS986449 | NC1=NC(C2=C(F)C([C@@H](C)N([C@H]3CCC(NC3=O)=O)C4=O)=C4C=C2)=CC(C)=C1C |
| BT33 | CC(C)C[C@@H]1CCO[C@]2([H])[C@@]([CN[C@]2([H])C(N[C@H]3[C@H](C)/C=C\C[C@H](F)CS[C@H]4O[C@@H]3[C@H](O[H])[C@H](O)[C@H]4O)=O)([H])C1 |
| Camibirstat | C[C@@H]1CN(C[C@@H](O1)C)C2=CC=CC(=N2)C3=CSC(=N3)NC(=O)[C@H](COC)NC(=O)C4=CN(C=C4)S(=O)(=O)C |
| CHK336 | O=C(C1=CSC(N2N=C(C(C3=CC=C(S(=O)(N)=O)C(F)=C3)=C2CC4CC4)C5=CC=C(F)C=C5)=N1)O |
| Compound36 | O=C(C(C)(C)C(F)F)N1C[C@H]2OC3=C(C#N)C=NC=C3[C@@H]1C2 |
| Emraclidine | CC1=CC(=NC2=C1CN(C2)C(=O)CC3CN(C3)C4=CC(=NC=C4)C(F)(F)F)C |
| Enozertinib | O=C(C=C)NC1=CC(NC2=NC=NC(N3OCC[C@@H]3C4=CC(F)=CC(F)=C4)=C2)=C(OC)C=C1N5CCC(N6CCN(C7CC7)CC6)CC5 |
| FG2101 | N#CC1CN(C1)C[C@H](C2=CC=C(C=C2)C#CC3=CC=C(C=C3)CN4CCOCC4)CC5=C(C(NC=N5)=O)OP(O)(O)=O |
| Fulacimstat | CN1C2=C(C=C(C=C2)N3C=C(C(=O)N(C3=O))[C@@H]4CCCC=C4C=CC=C5C(F)(F)F)C(=O)O)OC1=O |
| GS2278 | C[C@@H](OC(NC1=C(C2=NC=C(NC(C3=CN=C(C(F)(F)F)N=C3)=O)C=C2)N=NN1C)=O)C4=C(F)N=CC(F)=C4 |
| GSK4418959 | O=C(NC1=CC=C(C(F)(F)F)C=C1F)[C@@H]2CCC[C@H](C3=CC=C(N(C)S(=O)(C4=CC(N=C(C)N5C)=C5C=C4)=O)C=C3)[C@H]2C(O)=O |
| GSK484 | CN1C2=C(C=C(C=C2OC)C(=O)N3CCC(C(C3)N)O)N=C1C4=CC5=CC=CC=C5N4CC6CC6.C1 |
| HRO761 | O=C(NC1C2=NC(C3=CCOCC3)=NN2C(C(N4CCN(C(C5=NC=NC(C)=C5O)=O)CC4)=C1CC)=O)NC6=CC=C(C(F)(F)F)C=C6C1 |
| HRX215 | O=C(C1=NNC2=C1C=C(C3=CC=NC=C3)C=N2)C4=C(F)C(NS(CCC)(=O)=O)=C(F)C=C4 |
| JNJ6640 | FC1=C(OC2=CC(C)=NN2)N=C(N3C[C@H](C4=CC=NC=C4)CC3)N=C1 |
| JNT517 | FC(C=C(C=C1)OC(F)(F)F)=C1CNC(N(C2CC2)[C@H]3CN(CCC3)C(N)=O)=O |
| JNT-517 | FC(C=C(C=C1)OC(F)(F)F)=C1CNC(N(C2CC2)[C@H]3CN(CCC3)C(N)=O)=O |
| Mezigdomide | FC1=CC(C#N)=CC=C1N2CCN(CC3=CC=C(COC4=CC=CC5=C4CN([C@H]6CCC(NC6=O)=O)C5=O)C=C3)CC2 |
| MK8527 | OC[C@]1(C#C)[C@@H](O)C[C@H](N2C3=NC(C1)=NC(N)=C3C=C2)O1 |
| MRTX1133 | OC1=CC(C2=C(F)C3=NC(OC[C@]45C[C@@H](F)CN4CCC5)=NC(N6C[C@H]7N[C@H](CC7)C6)=C3C=N2)=C8C(C#C)=C(F)C=CC8=C1 |
| Muvalaplin | OC([C@H]([C@H]1CCNC1)CC2=CC(CN(CC3=CC(C[C@@H]([C@H]4CCNC4)C(O)=O)=CC=C3)CC5=CC(C[C@@H]([C@H]6CCNC6)C(O)=O)=CC=C5)=CC=C2)=O |
| NVPDF332 | C1CC(C2=C(C13CN4C=C(C=C(C4=N3)OC(F)F)C(F)(F)F)C=CN=C2C1)(F)F |
| NVPEDI048 | O=C(OC)C1=C(C1)C=CC(N(C)C(C2=CC3=C(C4=CC=C(C(NC)=O)C=C4)C=NN3C=C2)=O)=C1 |
| Rezatapopt | FC(F)(F)CN1C2=CC=CC(N[C@H]3[C@H](CN(CC3)C)F)=C2C=C1C#CCNC4=C(C=C(C=C4)C(NC)=O)OC |
| RP1664 | CC1=CC(NC2=NC(N(C3=C(C=C(C=C3F)S(C)=O)=O)F)C)=NC(C4=CN(C=N4)C)=C2C5CC5)=NN1 |
| Seltorexant | CC1=CC(=NC(=N1)N2C[C@@H]3CN(C[C@@H]3C2)C(=O)C4=C(C=CC=C4F)N5N=CC=N5)C |
| Sevasemten | O=C1C=CC(C2=CN=C(OC(F)(F)F)N=C2)=NN1CC3=CC(F)=CN=C3 |
| TAK756 | O=C(N)C1=CC=CC(C2=NOC(C3=CC(CN(C)C4=O)=C4C=C3O[C@@H]5CC[C@H](O)CC5)=C2)=C1 |
| Tavapadon | CC1=C(C=CC(=C1)OC2=C(C=CC=N2)C(F)(F)F)C3=C(C(=O)NC(=O)N3)C |
| TYRA300 | C[C@H](C1=C(C=NC=C1Cl)Cl)OC2=CC3=C(C=C2)NN=C3C4=CN=C(C=C4)N5CC6(C5)CN(C6)S(=O)(=O)C |
| VVD214 | O=C(N[C@@H](C1CC1)/C=C/S(C)(=O)=O)C2=CN=C(C(F)(F)C)N=C2OC3=CC=CC=C3 |
| Zongertinib | O=C(NC1CCN(CC1)C2=NC=C3N=CN=C(C3=N2)NC4=CC=C(C(C)=C4)OC5=CC(N=CN6C)=C6C=C5)C=C |

**Supplementary Table 4.** Predicted *vs* observed f_a_ (numerical and classifications) for the Prosilico validation data set using the 5 software. f_a_<0.3 (red); f_a_=0.3-0.8 (yellow); f_a_>0.8 (green).

| | PREDICTED $f_a$ | | | | | | | |
| --- | --- | --- | --- | --- | --- | --- | --- | --- |
|  | ANDROMEDA | PkCSM | Swiss ADME |  |  | ADMETlab 3.0 | DruMAP |  |
| OBSERVED $f_a$ | | | Abs model | Solubility | Total | | | Compound |
| ca 0 | 0,00 | 0,18 | Low | Poor | Low | <0,3 | <0,2 | AB15 |
| ca 0 | 0,00 | 0,00 | Failed | Failed | Failed | <0,3 | <0,2 | AB3 |
| ca 0 | 0,01 | 0,92 | Low |  | Low | >0,3 | <0,2 | AB6 |
| ca 0 (very low Papp) | 0,00 | 0,00 | Low | Poor | Low | >0,3 | <0,2 | AB8 |
| Very low | 0,00 | 0,00 | Low |  | Low | >0,3 | <0,2 | AB32 |
| Very low (ca 0) | 0,00 | 0,00 | Low | Insoluble | Low | <0,3 | <0,2 | AB35 |
| Very low (ca 0) | 0,00 | Failed | Low | Insoluble | Low | <0,3 | <0,2 | AB36 |
| 0,01 | 0,00 | 0,00 | Failed | Failed | Failed | <0,3 | <0,2 | AB2 |
| 0,10-0,16 | 0,00 | 0,04 | Failed | Failed | Failed | <0,3 | <0,2 | AB14 |
| 0,15-0,18 | 0,02 | 0,39 | High | Poor | Inconclusive | >0,3 | <0,2 | AB13 |
| 0,23-0,25 | 0,62 | 0,94 | Low |  | Low | >0,3 | <0,2 | AB25 |
| <0,3 | 0,38 | 0,93 | High | Poor | Inconclusive | <0,3 | <0,2 | AB41 |
| <0,3 | 0,54 | 0,81 | High |  | High | >0,3 | 0,2-0,7 | AB42 |
| 0,3 | 0,59 | 0,99 | Low | Insoluble | Low | >0,3 | 0,2-0,7 | AB44 |
| 0,3 | 0,71 | 1,00 | Low | Poor | Low | >0,3 | 0,2-0,7 | AB45 |
| 0,3 | 0,84 | 0,94 | Low | Poor | Low | >0,3 | Failed | AB46 |
| 0,3 | 0,16 | 0,89 | High | Poor | Inconclusive | >0,3 | >0,7 | AB47 |
| 0,3 | 0,58 | 0,95 | High |  | High | >0,3 | 0,2-0,7 | AB48 |
| 0,33 | 0,70 | 0,89 | Low |  | Low | >0,3 | >0,7 | AB11 |
| ca 1/3 | 0,58 | 0,98 | Failed | Failed | Failed | >0,3 | >0,7 | AB31 |
| ca 1/3 | 0,31 | 0,79 | Low | Insoluble | Low | >0,3 | >0,7 | AB39 |
| 0,35 | 0,61 | 1,00 | Low |  | Low | >0,3 | 0,2-0,7 | AB24 |
| 0,40 | 0,34 | 0,53 | Low |  | Low | >0,3 | 0,2-0,7 | AB12 |
| ca 0,4 | 0,68 | 0,94 | High | Poor | Inconclusive | >0,3 | >0,7 | AB38 |
| 0,5 | 0,81 | 0,94 | High | Poor | Inconclusive | >0,3 | >0,7 | AB1 |
| ca 0,5 | 0,62 | 0,85 | High | Poor | Inconclusive | >0,3 | 0,2-0,7 | AB30 |
| 0,50 | 0,47 | 0,93 | Failed | Failed | Failed | >0,3 | >0,7 | AB40 |
| 0,55 | 0,89 | 0,87 | High | Poor | Inconclusive | >0,3 | >0,7 | AB19 |
| 0,60 | 0,30 | 0,26 | Low | Poor | Low | >0,3 | 0,2-0,7 | AB4 |
| 0,70 | 0,43 | 0,91 | Low |  | Low | >0,3 | <0,2 | AB7 |
| >0,7-0,8 | 0,93 | 0,82 | High | Insoluble | Low | >0,3 | >0,7 | AB21 |
| >0,7-0,8 | 0,71 | 0,83 | High |  | High | >0,3 | >0,7 | AB37 |
| 0,81 | 0,57 | 0,93 | High | Poor | Inconclusive | >0,3 | 0,2-0,7 | AB18 |
| 0,85 | 0,83 | 0,93 | High |  | High | >0,3 | >0,7 | AB33 |
| 0,85 | 0,63 | 0,80 | High | Poor | Inconclusive | >0,3 | >0,7 | AB5 |
| 0,90 | 0,68 | 0,96 | High |  | High | >0,3 | >0,7 | AB16 |
| >0,9 | 0,86 | 0,77 | Low | Poor | Low | >0,3 | 0,2-0,7 | AB23 |
| 0,93 | 0,83 | 0,95 | High | Poor | Inconclusive | >0,3 | >0,7 | AB34 |
| >0,94 | 0,90 | 1,00 | High | Poor | Inconclusive | >0,3 | 0,2-0,7 | AB22 |
| ca 1 | 0,98 | 0,98 | High | Poor | Inconclusive | >0,3 | >0,7 | AB20 |
| ca 1 | 0,97 | 0,97 | High |  | High | >0,3 | >0,7 | AB50 |
| ca 1 (high Papp) | 1,00 | 1,00 | High | Poor | Inconclusive | >0,3 | >0,7 | AB10 |
| ca 1 (high Papp) | 1,00 | 0,96 | High | Insoluble | Low | >0,3 | >0,7 | AB9 |
| Near complete | 1,00 | 0,92 | High | Poor | Inconclusive | >0,3 | >0,7 | AB49 |

**Supplementary Table 5.** Predicted f_a_ (numerical and classifications) for the data set without available and published f_a_-values using the 5 software. f_a_<0.3 (red); f_a_=0.3-0.8 (yellow); f_a_>0.8 (green).

| PREDICTED $f_a$ | | | | | | | |
| --- | --- | --- | --- | --- | --- | --- | --- |
| ANDROMEDA | PkCSM | Swiss ADME<br>Abs model | Solubility | Total | ADMETlab 3.0 | DruMAP | Compound |
| 0,28 | 0,47 | Low |  | Low | >0,3 | 0,2-0,7 | BT33 |
| 0,35 | 0,85 | High | Poor | Inconclusive | >0,3 | 0,2-0,7 | Mezigdomide |
| 0,43 | 0,81 | High | Poor | Inconclusive | >0,3 | >0,7 | BMS986449 |
| 0,45 | 0,95 | High | Poor | Inconclusive | >0,3 | 0,2-0,7 | MRTX1133 |
| 0,46 | 0,08 | Low | Poor | Low | >0,3 | 0,2-0,7 | Muvalaplin |
| 0,47 | 1,00 | Low | Poor | Low | >0,3 | <0,2 | AMG410 |
| 0,49 | 0,70 | High |  | High | >0,3 | 0,2-0,7 | MK8527 |
| 0,50 | 0,73 | Low | Poor | Low | >0,3 | <0,2 | BAY3389934 |
| 0,50 | 0,74 | Low | Poor | Low | >0,3 | 0,2-0,7 | CHK336 |
| 0,51 | 0,67 | Low | Poor | Low | >0,3 | <0,2 | HRO761 |
| 0,52 | 0,69 | Low | Poor | Low | >0,3 | 0,2-0,7 | FG2101 |
| 0,55 | 0,72 | Low |  | Low | >0,3 | 0,2-0,7 | Camibirstat |
| 0,57 | 0,83 | High |  | High | >0,3 | >0,7 | AZD2389 |
| 0,57 | 0,96 | High |  | High | >0,3 | <0,2 | AZD0022 |
| 0,58 | 0,91 | High |  | High | >0,3 | >0,7 | JNT-517 |
| 0,59 | 0,77 | Low | Poor | Low | >0,3 | >0,7 | GS2278 |
| 0,60 | 1,00 | High | Poor | Inconclusive | >0,3 | 0,2-0,7 | Zongertinib |
| 0,60 | 0,95 | High |  | High | >0,3 | >0,7 | Compound36 |
| 0,62 | 0,74 | Low | Poor | Low | >0,3 | <0,2 | GSK4418959 |
| 0,63 | 0,64 | Low |  | Low | >0,3 | <0,2 | ALG000184 |
| 0,66 | 0,95 | High |  | High | >0,3 | >0,7 | Alixorexton |
| 0,66 | 0,85 | High | Poor | Inconclusive | >0,3 | 0,2-0,7 | Enozertinib |
| 0,67 | 1,00 | High | Insoluble | Low | >0,3 | 0,2-0,7 | BBO10203 |
| 0,68 | 0,85 | High | Poor | Inconclusive | >0,3 | >0,7 | AZD0780 |
| 0,68 | 0,89 | High | Poor | Inconclusive | >0,3 | 0,2-0,7 | TAK756 |
| 0,69 | 0,89 | High |  | High | >0,3 | 0,2-0,7 | GSK484 |
| 0,71 | 0,94 | High | Poor | Inconclusive | >0,3 | 0,2-0,7 | TYRA300 |
| 0,71 | 1,00 | High | Poor | Inconclusive | >0,3 | >0,7 | Sevasemten |
| 0,72 | 0,83 | Low | Poor | Low | >0,3 | 0,2-0,7 | HRX215 |
| 0,73 | 0,73 | High |  | High | >0,3 | 0,2-0,7 | Fulacimstat |
| 0,73 | 0,94 | High | Poor | Inconclusive | >0,3 | >0,7 | NVPED1048 |
| 0,75 | 0,87 | Low | Poor | Low | >0,3 | >0,7 | NVPDF332 |
| 0,76 | 0,95 | High |  | High | >0,3 | >0,7 | Emraclidine |
| 0,78 | 0,80 | Low | Poor | Low | >0,3 | 0,2-0,7 | RP1664 |
| 0,78 | 0,99 | High | Poor | Inconclusive | >0,3 | >0,7 | BI9508 |
| 0,81 | 0,94 | High | Poor | Inconclusive | >0,3 | >0,7 | Azetukalner |
| 0,81 | 0,88 | High | Poor | Inconclusive | >0,3 | 0,2-0,7 | Rezatapopt |
| 0,81 | 0,89 | Low | Poor | Low | >0,3 | >0,7 | ARN23746 |
| 0,86 | 0,96 | High |  | High | >0,3 | >0,7 | JNJ6640 |
| 0,88 | 0,95 | High | Poor | Inconclusive | >0,3 | 0,2-0,7 | Tavapadon |
| 0,97 | 0,83 | High |  | High | >0,3 | >0,7 | VVD214 |
| 0,97 | 0,99 | High |  | High | >0,3 | >0,7 | Seltorexant |

**Supplementary Table 6.** Predicted *vs* observed f_u_ (numerical and classifications) for the Prosilico validation data set using 4 of the 5 software. f_u_<0.01 (red); f_u_=0.01-0.03 (orange); f_u_=0.03-0.1 (light yellow); f_u_=0.1-0.3 (yellow); f_u_>0.6 (blue); f_u_>0.6 (green).

| | PREDICTED $f_u$ | | | | | |
| --- | --- | --- | --- | --- | --- | --- |
| OBSERVED $f_u$ | ANDROMEDA | PkCSM | Swiss ADME | ADMETlab 3.0 | DruMAP | Compound |
| <0,001 | 0,0006 | Failed (0) | n.a. | 0,007 | 0,009 | D1 |
| <0,001 | 0,0067 | 0,34 | n.a. | 0,003 | 0,003 | D72 |
| <0,001 | 0,0037 | 0,37 | n.a. | 0,005 | 0,013 | D73 |
| 0,001 | 0,0025 | 0,026 | n.a. | 0,006 | 0,008 | D10 |
| <0,003 | 0,0016 | 0,29 | n.a. | 0,016 | 0,026 | D26 |
| 0,004 | 0,0027 | 0,25 | n.a. | 0,014 | 0,006 | D50 |
| <0,005 | 0,0067 | 0,37 | n.a. | 0,012 | 0,010 | D78 |
| 0,005 | 0,0196 | Failed (0) | n.a. | 0,015 | 0,010 | D2 |
| 0,005 | 0,0046 | 0,26 | n.a. | 0,007 | 0,011 | D11 |
| 0,005 | 0,0045 | 0,15 | n.a. | 0,007 | 0,006 | D13 |
| 0,005 | 0,0079 | 0,070 | n.a. | 0,007 | 0,005 | D20 |
| 0,005 | 0,005 | 0,083 | n.a. | 0,010 | 0,023 | D27 |
| 0,0061 | 0,0099 | 0,16 | n.a. | 0,013 | 0,006 | D48 |
| 0,008 | 0,0058 | 0,034 | n.a. | 0,008 | 0,004 | D52 |
| <0,01 | 0,0064 | 0,029 | n.a. | 0,011 | 0,002 | D66 |
| <0,01 | 0,036 | 0,14 | n.a. | 0,020 | 0,027 | D68 |
| <0,01 | 0,072 | 0,060 | n.a. | 0,007 | 0,005 | D75 |
| <0,01 | 0,015 | Failed (0) | n.a. | 0,050 | 0,096 | D76 |
| <0,01 | 0,003 | 0,071 | n.a. | 0,205 | 0,080 | D12 |
| <0,01 | 0,0024 | Failed (0) | n.a. | 0,008 | 0,001 | D14 |
| <0,01 | 0,0063 | 0,054 | n.a. | 0,009 | 0,016 | D25 |
| <0,01 | 0,004 | Failed (0) | n.a. | 0,009 | 0,004 | D28 |
| <0,01 | 0,019 | 0,10 | n.a. | 0,026 | 0,012 | D39 |
| 0,01 | 0,013 | 0,13 | n.a. | 0,024 | 0,016 | D5 |
| 0,01 | 0,063 | 0,10 | n.a. | 0,025 | 0,024 | D41 |
| ca 0,01 | 0,013 | Failed (0) | n.a. | 0,32 | 0,044 | D9 |
| 0,011 | 0,0089 | 0,011 | n.a. | 0,005 | 0,006 | D18 |
| <0,012 | 0,0097 | 0,14 | n.a. | 0,008 | 0,018 | D74 |
| 0,015 | 0,008 | Failed (0) | n.a. | 0,014 | 0,015 | D23 |
| <0,02 | 0,021 | Failed (0) | n.a. | 0,012 | 0,047 | D69 |
| <0,02 | 0,019 | 0,094 | n.a. | 0,017 | 0,016 | D70 |
| 0,023 | 0,017 | 0,23 | n.a. | 0,042 | 0,066 | D42 |
| 0,025 | 0,079 | 0,18 | n.a. | 0,064 | 0,017 | D71 |
| 0,0275 | 0,017 | 0,12 | n.a. | 0,011 | 0,016 | D51 |
| 0,030 | 0,020 | 0,25 | n.a. | 0,028 | 0,234 | D43 |
| 0,04 | 0,019 | 0,20 | n.a. | 0,15 | 0,029 | D38 |
| 0,05 | 0,025 | 0,15 | n.a. | 0,074 | 0,018 | D15 |
| 0,1 | 0,062 | 0,062 | n.a. | 0,077 | 0,062 | D40 |
| 0,17 | 0,12 | 0,25 | n.a. | 0,052 | 0,044 | D44 |
| 0,43 | 0,47 | 0,26 | n.a. | 0,43 | 0,070 | D8 |
| 0,45 | 0,19 | 0,25 | n.a. | 0,49 | 0,53 | D4 |
| 0,54 | 0,46 | 0,41 | n.a. | 0,23 | 0,48 | D46 |
| 0,61 | 0,40 | 0,38 | n.a. | 0,25 | 0,37 | D49 |
| 0,74 | 0,51 | 0,51 | n.a. | 0,76 | 0,22 | D37 |
| 0,75 | 0,63 | 0,52 | n.a. | 0,70 | 0,55 | D6 |
| 0,76 | 0,86 | 0,80 | n.a. | 0,82 | 0,73 | D17 |
| 0,77 | 0,73 | 0,54 | n.a. | 0,72 | 0,39 | D19 |
| 0,77 | 0,37 | 0,39 | n.a. | 0,39 | 0,62 | D3 |
| 0,85 | 0,85 | 0,75 | n.a. | 0,87 | 0,34 | D24 |
| 0,89 | 0,79 | 0,18 | n.a. | 0,024 | 0,030 | D45 |
| 0,90 | 0,85 | 0,70 | n.a. | 0,87 | 0,69 | D16 |
| 0,92 | 0,86 | 0,74 | n.a. | 0,65 | 0,63 | D7 |
| 0,98 | 0,97 | 0,45 | n.a. | 0,43 | 0,74 | D21 |
| 1,00 | 0,71 | 0,42 | n.a. | 1,00 | 0,60 | D47 |

**Supplementary Table 7.** Predicted f_u_ (numerical and classifications) for the data set without available and published f_u_-values using 4 of the 5 software. f_u_<0.01 (red); f_u_=0.01-0.03 (orange); f_u_=0.03-0.1 (light yellow); f_u_=0.1-0.3 (yellow); f_u_>0.6 (blue); f_u_>0.6 (green).

| ANDROMEDA | PkCSM | Swiss ADME | ADMETlab 3.0 | DruMAP |  |
| --- | --- | --- | --- | --- | --- |
| 0,001 | 0,210 | n a | 0,014 | 0,012 | GSK4418959 |
| 0,004 | 0,211 | n a | 0,007 | 0,005 | CHK336 |
| 0,005 | 0,149 | n a | 0,032 | 0,042 | BBO10203 |
| 0,011 | No/Zero fu | n a | 0,021 | 0,027 | HRO761 |
| 0,014 | 0,189 | n a | 0,020 | 0,006 | NVPED1048 |
| 0,016 | 0,206 | n a | 0,027 | 0,043 | Zongertinib |
| 0,021 | 0,055 | n a | 0,014 | 0,036 | TYRA300 |
| 0,022 | No/Zero fu | n a | 0,018 | 0,007 | Azetukalner |
| 0,023 | 0,103 | n a | 0,068 | 0,037 | GS2278 |
| 0,023 | 0,153 | n a | 0,038 | 0,038 | RP1664 |
| 0,028 | 0,137 | n a | 0,009 | 0,015 | Tavapadon |
| 0,035 | 0,160 | n a | 0,013 | 0,020 | Fulacimstat |
| 0,037 | 0,149 | n a | 0,023 | 0,019 | BI9508 |
| 0,043 | 0,146 | n a | 0,068 | 0,034 | ARN23746 |
| 0,044 | 0,244 | n a | 0,018 | 0,123 | Enozertinib |
| 0,050 | 0,116 | n a | 0,027 | 0,037 | Rezatapopt |
| 0,050 | 0,058 | n a | 0,023 | 0,013 | HRX215 |
| 0,055 | 0,028 | n a | 0,040 | 0,042 | Mezigdomide |
| 0,055 | No/Zero fu | n a | 0,019 | 0,004 | VVD214 |
| 0,063 | 0,176 | n a | 0,092 | 0,006 | NVPDF332 |
| 0,067 | 0,224 | n a | 0,029 | 0,024 | BAY3389934 |
| 0,072 | 0,305 | n a | 0,137 | 0,080 | AZD0780 |
| 0,079 | 0,077 | n a | 0,044 | 0,033 | Seltorexant |
| 0,087 | 0,204 | n a | 0,047 | 0,035 | Emraclidine |
| 0,099 | 0,308 | n a | 0,040 | 0,021 | ALG000184 |
| 0,103 | 0,276 | n a | 0,154 | 0,029 | AZD0022 |
| 0,106 | 0,134 | n a | 0,060 | 0,029 | Camibirstat |
| 0,108 | 0,164 | n a | 0,035 | 0,040 | AMG410 |
| 0,112 | 0,265 | n a | 0,177 | 0,163 | Muvalaplin |
| 0,123 | 0,131 | n a | 0,367 | 0,044 | TAK756 |
| 0,125 | 0,275 | n a | 0,247 | 0,103 | Sevasemten |
| 0,133 | 0,265 | n a | 0,073 | 0,023 | JNJ6640 |
| 0,136 | 0,098 | n a | 0,066 | 0,158 | GSK484 |
| 0,137 | 0,321 | n a | 0,163 | 0,029 | MRTX1133 |
| 0,149 | 0,148 | n a | 0,136 | 0,278 | BMS986449 |
| 0,247 | 0,351 | n a | 0,074 | 0,193 | Compound36 |
| 0,247 | 0,249 | n a | 0,093 | 0,018 | FG2101 |
| 0,267 | 0,268 | n a | 0,039 | 0,119 | Alixorexton |
| 0,336 | 0,361 | n a | 0,029 | 0,045 | JNT-517 |
| 0,355 | 0,530 | n a | 0,152 | 0,131 | BT33 |
| 0,396 | 0,527 | n a | 0,580 | 0,604 | MK8527 |
| 0,445 | 0,211 | n a | 0,600 | 0,139 | AZD2389 |

